# Comparative characterization of Cas12a2 orthologs identifies high-activity nucleases for programmable cell elimination

**DOI:** 10.64898/2026.06.23.734040

**Authors:** Anna L. Singer, Emma E. January, Erin K. Zess, Allison J. N. Antonakos, Matthew B. Begemann

## Abstract

Cas12a2 CRISPR nucleases, including SuCas12a2, have been shown to have extensive collateral activity towards RNA, ssDNA, and dsDNA. This collateral activity results in targeted cell elimination and has applications across biotechnology, agriculture, and human health. We explored the natural genetic diversity of Cas12a2 nucleases and characterized nine novel orthologs in a DNA damage kinetic assay in E. coli. Three new Cas12a2 orthologs (RsCas12a2, SdCas12a2, and HmCas12a2) were shown to have high collateral activity towards DNA. These nucleases are highly divergent from SuCas12a2, have conserved core RuvC catalytic residues, and have sequence diversity in the previously reported aromatic clamp residues required for nucleic acid positioning in the active site. We defined PFS preferences and mismatch tolerance for each high-activity Cas12a2 nuclease, expanding the available Cas12a2 toolbox, and discovered functional differences with obvious impacts on downstream applications.

## Introduction

CRISPR–Cas systems are RNA-guided, nucleic-acid-targeting proteins used by bacteria and archaea to defend against bacteriophages, plasmids, and other foreign genetic elements [1,2]. Beyond their native role in adaptive immunity, members of the broader Cas nuclease family have been applied as genome engineering and diagnostic tools, including precision genome editing (Cas9, Cas12a) [3,4] and transcript-level detection (Cas13) [5].

Multiple CRISPR–Cas systems have been shown to have collateral activity, also known as *trans*-cleavage activity, resulting in the indiscriminate degradation of nucleic acid substrates following initial target-mediated activation [6,7]. This secondary activity has either been viewed as an off-target effect, such as with Cas12a degradation of ssDNA [6], or as a usable feature to target the degradation of nucleic acids, such as with Cas13 collateral activity against RNA [7]. Cas12a2 is a group of Type V CRISPR nuclease proteins that use non-specific collateral activity as a main mode of action in defense. Cas12a2 collateral activation occurs after RNA-guided recognition and results in non-specific cleavage of RNA, ssDNA, and dsDNA. This large-scale, indiscriminate nucleic acid damage triggers an SOS-driven DNA damage response in the bacterial host that leads to cell elimination via apoptosis [8]. The biochemical and structural basis of Cas12a2 collateral activity has been determined for SuCas12a2 (from *Sulfuricurvum* sp. PC08-66), showing that RNA-target binding restructures the RuvC catalytic site to allow for non-specific accommodation of dsDNA, ssDNA, and ssRNA substrates [9].

The unique collateral activity of Cas12a2 has been leveraged across research areas. SuCas12a2 has been used in diagnostics for sensitive detection of SARS-CoV-2 [10], as well as *Entamoeba histolytica* and *Mycoplasma pneumoniae* infection [11]. SuCas12a2 and the related nuclease GeCas12a2 have been demonstrated to perform targeted cell elimination in *S. cerevisiae* and multiple human cell lines [12]. Newly published work quantified SuCas12a2-mediated DNA damage in human cells and harnessed subsequent DNA damage-induced apoptosis for the selective elimination of cells harboring oncogenic mutations [12,13]. This research also explored target transcript abundance, targeting specificity, and mouse model delivery and collectively suggests that SuCas12a2 could be a viable gene therapy for specific oncology applications, including the “undruggable” p53 oncogenic mutations [13]. The diagnostic capabilities of Cas12a2 have also been combined with oncology applications, with a recent publication showing that Cas12a2 can be turned into a tumor visualization nanoprobe that binds to oncogenic mRNA [14].

The potential impact across broad applications such as agriculture, diagnostics, and human health led us to explore the natural diversity of Cas12a2 nucleases to identify novel nucleases with unique features relative to SuCas12a2, including kinetic activity, specificity, and targeting ability. We identified SuCas12a2 orthologs from publicly available genomic data and used a preliminary screen to identify functional Cas12a2 variants. We characterized the DNA damage kinetics of nine new Cas12a2 orthologs and described the protospacer flanking sequence (PFS) preferences and target mismatch tolerance. We identified three high-activity nucleases for programmable cell elimination with unique sequence features and functional capabilities.

## Results

### Functional characterization of Cas12a2 orthologs identifies three high-activity novel nucleases

We mined public genomic data for full-length Cas12a2 sequences and identified SuCas12a2 orthologs from a diverse range of host-associated and environmental metagenomes including *Reticulitermes speratus*, *Procryptotermes leewardensis*, *Hodotermes mossambicus*, and *Euhamitermes* sp. (all termite-gut), *Sus domesticus* (pig gut), peat permafrost (Sweden), mine drainage, and two landfill metagenomes (Figure 1a, Supplementary Figure 1). Each ortholog was named with a 2-letter prefix derived from its bacterial source organism or, where the bacterium is unclassified, from its eukaryotic host. The complete inventory of source organisms, NCBI protein accessions, and corresponding internal designations for each ortholog is provided in Supplementary Table S1.

**Figure 1.**
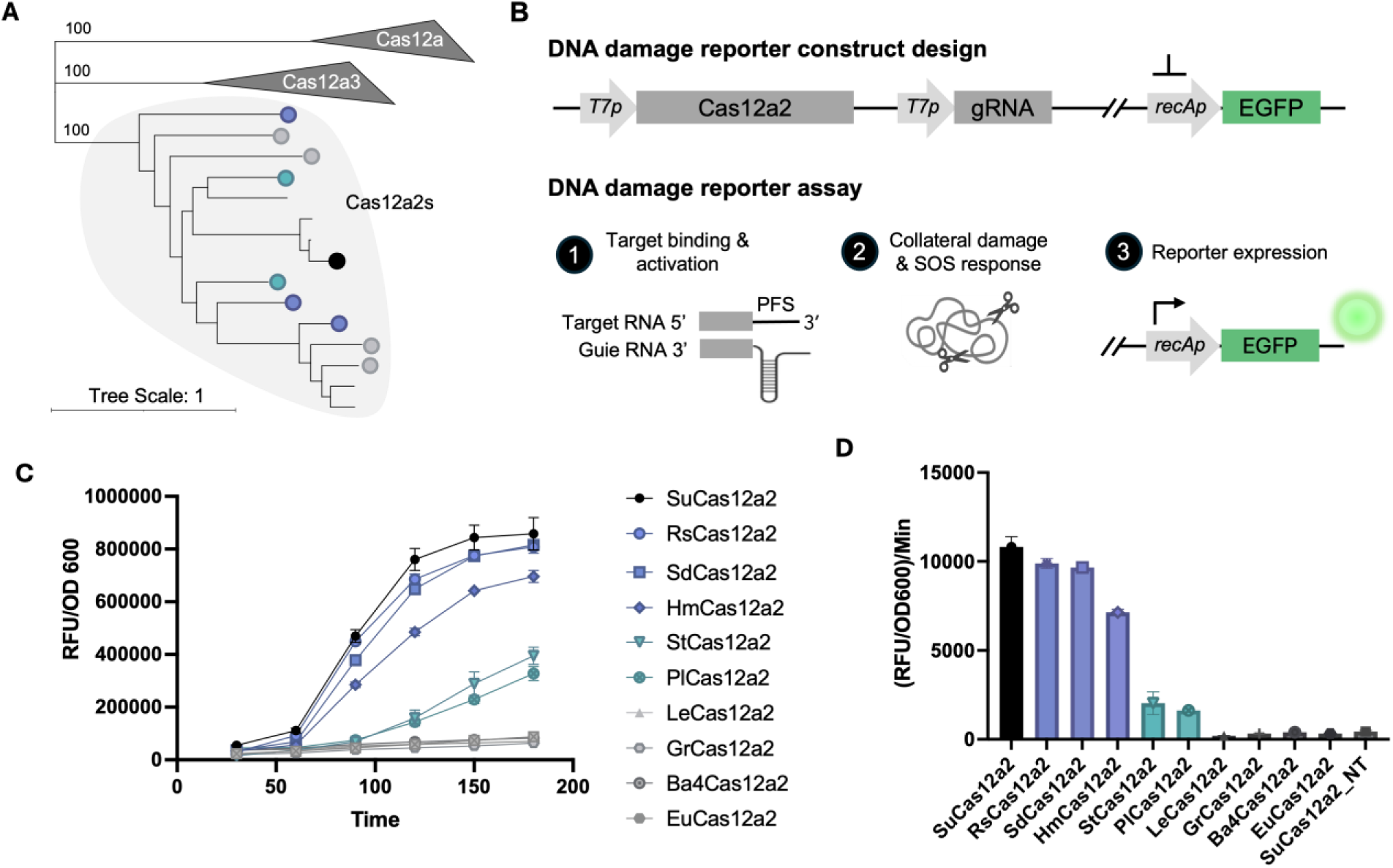
Diverse Cas12a2 orthologs have varying DNA damage activity. (A) Maximum likelihood phylogenetic tree of 24 Cas12 nucleases, with outgroup Cas12a and Cas12a3 clades collapsed (Supplementary Figure 1). Coloring corresponds to activity designations based on functional characterization. (B) Schematic of the DNA damage kinetic assay. Reporter construct includes inducible Cas12a2 and gRNA expression cassettes in addition to a GFP reporter driven by the *recA* promoter. Target binding activates collateral activity triggering SOS response and subsequent reporter expression. (C) Representative *recA*-GFP reporter kinetic curves for 9 novel nucleases and SuCas12a2. (D) Slopes of *recA*-GFP reporter S-curves calculated in best fitting linear range, benchmarked to SuCas12a2 and arranged by activity groupings.

To rapidly screen the Cas12a2 orthologs for collateral activity, we developed a DNA damage kinetic assay in *Escherichia coli* that quantifies target-dependent DNA damage (Figure 1b). This assay measures GFP expression driven by the *recA* promoter, which is induced in response to DNA damage, with fluorescence normalized to cell growth (RFU/OD₆₀₀) (Figure 1b). These measurements were taken every 30 minutes over a 3-hour growth period to establish linear ranges and then in-range data points are used to calculate the rate of change of activity (Figure 1c). The assay uses a gRNA and target sequence previously shown to have high activity across Cas12a and Cas12a2 nucleases [15,8], and on-target activity was compared with a paired non-target control (Supplementary Figure 2).

We observed that three of the nine new Cas12a2 orthologs—SdCas12a2, RsCas12a2, and HmCas12a2—had DNA damage activity on par with SuCas12a2 (Figure 1d). Two other orthologs, StCas12a2 and PlCas12a2, had intermediate DNA damage activity (Figure 1d). The remaining four orthologs—LeCas12a2, Ba4Cas12a2, EuCas12a2, and GrCas12a2—produced DNA damage activity within range of the non-target control and were classified as having background levels of activity (Figure 1d). Interestingly, Cas12a2 nucleases with differing DNA damage activity levels are found within the same phylogenetic subclades (Figure 1a, Supplementary Figure 1), suggesting multiple evolutionary origin points of high DNA damage activity.

Despite the background-level performance of LeCas12a2, Ba4Cas12a2, EuCas12a2, and GrCas12a2 in the DNA damage kinetic assay, all nine new Cas12a2 orthologs showed collateral activity in a plate-based bacterial toxicity assay (Supplementary Figure 3). This finding suggests that the Cas12a2 background-level activity group may be slower to accumulate DNA damage or may have an alternative collateral activity mode of action that is not captured by the DNA damage kinetic assay. Overall, the initial findings suggest that sequence-relatedness is insufficient to predict Cas12a2 nucleases with high DNA damage activity (Figure 1).

### High-activity Cas12a2 orthologs are sequence-diverse with a conserved catalytic core

To better understand the biochemical basis of differential Cas12a2 DNA damage activity, we compared amino acid sequence conservation across the Cas12a2 orthologs. This analysis showed that the Cas12a2 orthologs have high levels of sequence divergence, with pockets of high conservation and low gap percentages concentrated around key functional residues (Figure 2a). Pairwise amino acid sequence identity comparisons of the high-activity Cas12a2 orthologs and SuCas12a2, show that RsCas12a2, SdCas12a2, and HmCas12a2 retain only 37–49% identity to SuCas12a2, as well as to one another (Figure 2b). In contrast, the recently characterized nuclease GeCas12a2 [12] is highly related to SuCas12a2 (Supplementary Figure 1).

**Figure 2.**
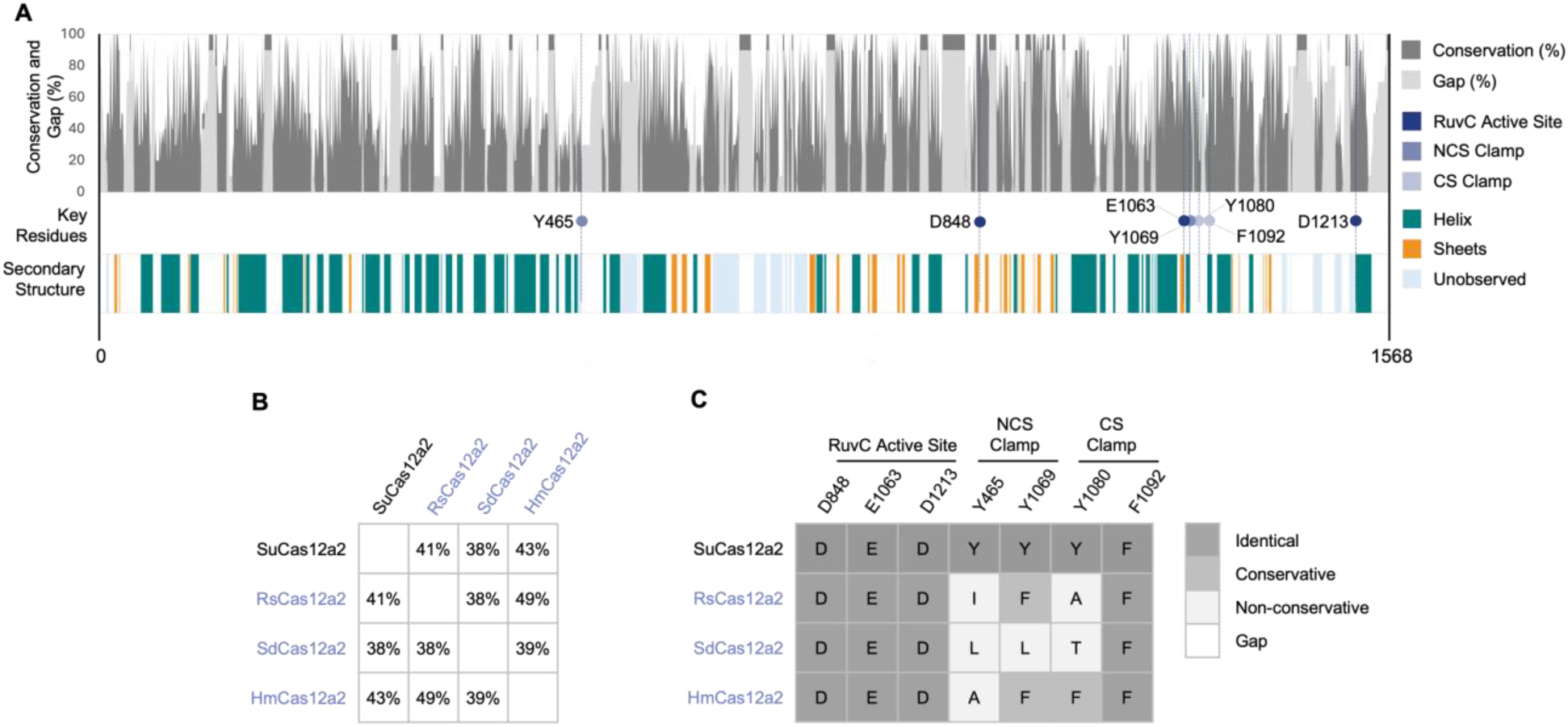
High-activity Cas12a2 orthologs are sequence-diverse with a conserved catalytic core. (A) Sequence conservation (non-gap) (%) and gap (%) (top) of the ten Cas12a2 sequences functionally characterized in this publication. Secondary structure (bottom) and key residues (middle) mapped from the SuCas12a2 binary crystal structure (PDB 8D49) [9]. (B) Percent identity matrix of the full-length sequences and (C) key residue mapping of SuCas12a2 and high-activity Cas12a2 nucleases (blue). Amino acid conservation at each site was scored as identical, conservative (preserves the chemical property), or non-conservative (major chemical change) relative to SuCas12a2.

Despite the full-length sequence divergence of the Cas12a2 orthologs, the catalytic core is conserved across all tested nucleases (Supplementary Figure 4). Each of the three RuvC active site residues (D848, E1063, D1213) are identical in the high-activity nucleases (SdCas12a2, RsCas12a2, HmCas12a2) and SuCas12a2 (Figure 2c). The cleavage strand (CS) aromatic clamp residue F1092 is similarly conserved, but the other key functional residues that form the aromatic clamp involved in nucleic acid positioning are more variable (Figure 2c). The CS aromatic clamp residue (Y1080) and the two non-cleavage strand (NCS) aromatic clamp residues (Y465 and Y1069) have conservative and non-conservative substitutions relative to SuCas12a2 in the high-activity nucleases (Figure 2c) and other tested nucleases (Supplementary Figure 4). The observed diversity within nucleic acid positioning residues led us to explore whether high-activity Cas12a2 orthologs show unique preferences for protospacer flanking sequence (PFS) recognition and mismatch tolerance.

### Protospacer flanking sequence (PFS) and mismatch profiling reveals distinct recognition preferences across the high-activity Cas12a2 orthologs

To characterize the protospacer flanking sequence (PFS) preferences of the nucleases, we tested the three high-activity Cas12a2 orthologs (RsCas12a2, SdCas12a2, HmCas12a2) and SuCas12a2 against seven 3′-PFS variants using the DNA damage kinetic assay. The seven PFS variants were selected based on results of a full library PFS characterization previously published for SuCas12a2 [8]. Selections included the canonical AAAG PFS, single substitutions at positions -2, -3, and -4 (AGAG, GAAG, AAGG), and progressively more divergent sequences (ACUU, GUGG, and ACCA) (Figure 3a). The absolute performance of each Cas12a2 ortholog at the canonical PFS was assessed and used to normalize activity at non-canonical PFS sequences (Figure 3b).

**Figure 3.**
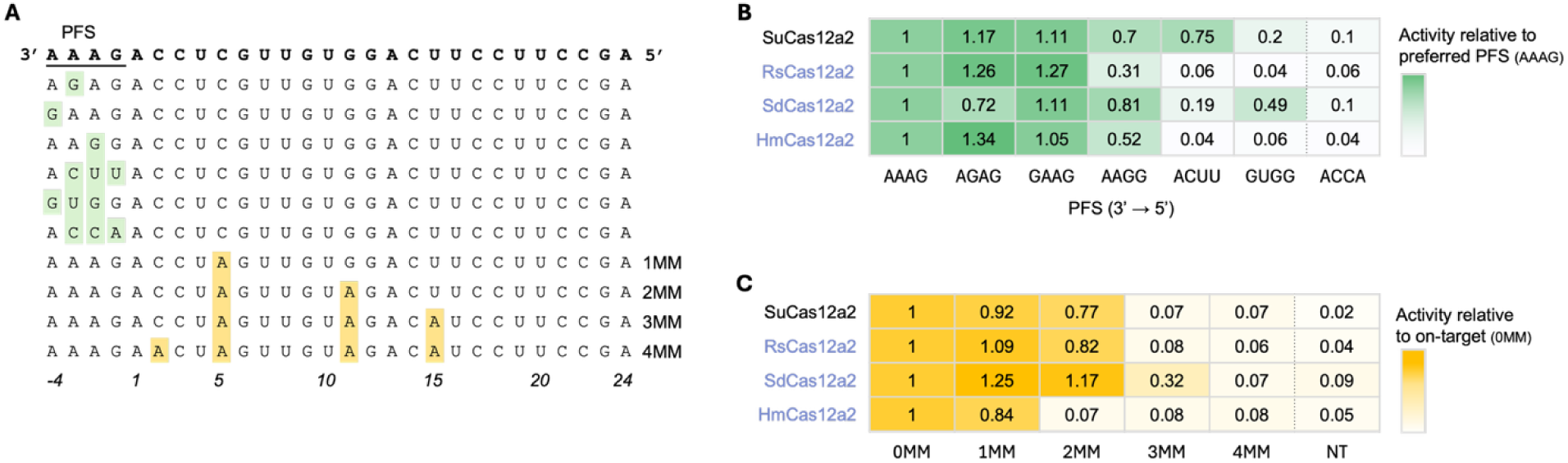
High-activity Cas12a2 orthologs have distinct PFS profiles and mismatch tolerance. (A) Cumulative PFS profile and target mismatch series tested in this work using the DNA damage kinetic assay. Top-most sequence represents the canonical PFS (AAAG) and perfect matching RNA target used to benchmark activity across all variants. PFS variants are listed with base substitutions highlighted in green. Mismatch variants are listed with base substitutions highlighted in yellow. (B) Heat map of PFS variant activity relative to AAAG canonical PFS. (C) Heat map of mismatch variant activity relative to 0MM target.

SuCas12a2 retained substantial DNA damage activity at AGAG, GAAG, AAGG, and ACUU, broadly consistent with its previously reported tolerance for non-canonical PFS sequences [8]. Interestingly, RsCas12a2 showed a narrower range of tolerated PFS sequences and produced higher relative activity at AGAG and GAAG than at the canonical AAAG PFS sequence (*p* value < 0.005). SdCas12a2 displayed a uniquely nuanced PFS preference, showing moderate DNA damage activity at the previously inaccessible GUGG PFS and robust DNA damage activity across AGAG, GAAG, and AAGG (Figure 3b). These diverse PFS preferences could allow RsCas12a2 and SdCas12a2 to be deployed for specific applications beyond what is currently achievable with SuCas12a2.

We next explored whether the top three high-activity Cas12a2 orthologs would also show divergent tolerance for mismatched base pairs within the targeting sequence. We synthesized a series of targeting plasmids containing 1-, 2-, 3-, or 4-mismatched base pairs with our guide RNA (Figure 3a). Activity for each target was normalized to the no-mismatch control, and DNA damage activity was assessed using the DNA damage kinetic assay. SdCas12a2 showed the most tolerance for mismatches with its protospacer, producing strong to modest DNA damage activity in the 1-, 2-, and 3-mismatch target variants. SuCas12a2 and RsCas12a2 show a similar tolerance profile, producing DNA damage activity at -1 and 2-mismatch targets. HmCas12a2 demonstrates the most stringency of the set, showing background levels of activity at targets with more than 1 mismatch (Figure 3c), making this nuclease a strong candidate to target transcript sequences with closely related off-targets.

## Discussion

Here we have functionally characterized nine new Cas12a2 orthologs and identified three nucleases with high DNA damage activity—SdCas12a2, RsCas12a2, and HmCas12a2 (Figure 1). The RuvC active site residues responsible for nucleic acid cleavage are highly conserved across all ten tested Cas12a2 orthologs, suggesting, unsurprisingly, that these catalytic sites are evolutionarily constrained (Figure 2). In contrast, the aromatic clamp residues are more divergent across the Cas12a2 orthologs (Figure 2). For SuCas12a2, these aromatic clamp residues have been shown to be responsible for positioning the nucleic acid substrate within the active site and are critical for collateral activity [9]. The lack of conservation in the Cas12a2 orthologs suggests that these nucleases may undergo different conformational changes upon initial target cleavage to accommodate nucleic acid positioning within the active site and enable collateral activity. Future work will need to address how the sequence differences of functional Cas12a2 family members influence protein folding dynamics and substrate engagement.

The protospacer flanking sequence (PFS) is a key feature of Cas12a2 specificity. The four base-pair sequence, which can vary from the canonical sequence (3′ AAAG 5′), must be present immediately downstream of the protospacer on the target transcript for Cas12a2 to function. Specifically, recognition of the PFS triggers Cas12a2 to undergo a conformational change that exposes the RuvC active site for initial RNA cleavage and subsequent collateral activity of dsDNA, ssDNA, and ssRNA [8,9]. In this study, we identified high-activity Cas12a2 nucleases with both broader and narrower PFS preferences relative to SuCas12a2 (Figure 3b), providing new tools to access more target sites or enhance specificity, respectively. In the former case, certain oncogenic mutations do not have a canonical PFS (3′ AAAG 5′) within range. In this situation, such as with the S37 mutation in CTNNB1, part of the exon 3 hotspot for hepatocellular carcinoma [27], SdCas12a2 may be able to access this target site because it can recognize a G-rich PFS, as seen in our panel (GUGG).

Off-target activation is of particular importance for potential oncological applications of Cas12a2. Most oncogenic mutations are patient-specific, single base pair mutations in crucial genes [23,24]. Cas12a2-activated DNA damage must be specific enough to eliminate cells with single base pair resolution while leaving normal, healthy cells intact. Cas12a2 specificity will rely on a combination of PFS suitability and protospacer mismatches to determine the likelihood of any unwanted Cas12a2 collateral activity. On the side of greater PFS specificity, we identified RsCas12a2 which has a narrow preference for only three tested PFS sequences (Figure 3b). This nuclease may better fit human health applications, where it is important to limit off-target activity and broad PFS access can lead to more off-target sites [22]. In this work, we have also shown that HmCas12a2 exhibits DNA damage activity that is less tolerant of mismatches within the protospacer as compared to other members of the Cas12a2 family when tested in our DNA damage kinetic assay. SuCas12a2 has been previously shown to have single base pair specificity in mammalian cell culture systems [13,12], suggesting that the *E. coli*-based kinetics assay used here has a higher level of sensitivity for DNA damage or that the physiological conditions of a mammalian cell culture system are sufficiently different to limit mismatch activity. By comparing the results of SuCas12a2 in mammalian culture systems with the results presented here, we posit that HmCas12a2 would have similar or better mismatch tolerance to SuCas12a2 in a eukaryotic system.

Together, RsCas12a2, SdCas12a2, and HmCas12a2 provide complementary additions to the Cas12a2 toolkit, with each ortholog suited to different application spaces. We have shown that these new high-activity Cas12a2 orthologs are not interchangeable with SuCas12a2, and each offers distinct features in activity, PFS recognition, and mismatch tolerance that could be selected for specific therapeutic or industrial applications. Collectively, the work presented here expands our understanding of Cas12a2 functional diversity, demonstrating that meaningful differences in nuclease behavior can be found across family members. As the Cas12a2 family continues to be explored, key principles will need to be established to help guide the selection and development of Cas12a2 tools with robust nucleic acid damage activity and high specificity.

## Materials and Methods

### Bacterial Strain Development

Strains were developed in *E. coli* BL21-AI sourced from Invitrogen. BL21-AI cells were transformed using pACYC expressing Cas12a2 and guide RNA cassettes under T7 promoter and Lac repressor or T7 promoter, respectively. Strains developed for testing in the DNA damage kinetic assay also included a *recA-*driven GFP cassette on this plasmid. A second pBR322 plasmid carrying a 500–600 base pair target gene sequence, with T7 promoter or without, was transformed into strains already carrying the Cas12a2 and guide. Strains were grown under dual selection with kanamycin and chloramphenicol antibiotics and used for subsequent bacterial toxicity and DNA damage kinetic assays.

### Bacterial Assays

#### Plate-based Bacterial Toxicity Assay

Novel Cas12a2 nucleases were initially validated in *E. coli* using a bacterial toxicity assay. Dual plasmid strains were grown overnight in 5 mL cultures of LB broth with selection. The next day, 3 replicates of fresh cultures were inoculated to a normalized OD around 0.02 and allowed to incubate for 2 hours at 37°C while shaking at 250 rpm. Expression of the Cas12a2 and guide was then induced by addition of 5 μL 1 M IPTG and 50 μL of 20% arabinose. After 2 hours of induction, a 10-fold dilution series was performed before plating on LB agar plates with chloramphenicol, maintaining selection for the Cas12a2 and gRNA plasmid. Colonies were counted the following day and survival of each treatment was recorded as a percent reduction compared to the non-target control.

#### DNA Damage Kinetic Assay

Strains were grown overnight in 5 mL cultures of LB with selection. The next day, 3 replicates of each treatment were inoculated into fresh 5 mL cultures to a normalized OD of around 0.02 and allowed to incubate for 2 hours at 37°C while shaking at 250 rpm. 100 μL of each treatment in triplicate were transferred into a clear-bottom 96-well plate in equal volume LB and induced with 2 μL 20% arabinose and 2 μL 0.1 M IPTG. OD₆₀₀ and GFP fluorescence were measured every 30 minutes over a 3-hour time course. Background-subtracted RFU/OD₆₀₀ was averaged over the 3 replicates at each time point to generate kinetic curves.

### Cas12a2 Sequence Identification

Cas12a2 sequences were identified through a BLAST analysis of available prokaryotic genome and metagenome sequences from NCBI and JGI using either the full-length protein sequence of SuCas12a2 or highly conserved functional domains.

### Phylogenetic Analysis

A maximum-likelihood phylogenetic tree of 15 Cas12a2 nucleases was calculated using MEGA12 [16] from a trimmed (ClipKIT [17]) amino acid sequence alignment (MUSCLE [18], codon-based) with Cas12a and Cas12a3 outgroups. The tree was visualized using iTOL [19] and bootstrap values were based on 1000 iterations.

### Sequence Alignment and Residue Mapping

A full-length sequence alignment (MUSCLE [18], codon-based) was performed with the ten characterized Cas12a2 nucleases (SuCas12a2, RsCas12a2, SdCas12a2, HmCas12a2, StCas12a2, PlCas12a2, Ba4Cas12a2, GrCas12a2, LeCas12a2, EuCas12a2) and the reference sequence GeCas12a2 [12]. The secondary structure elements and key residues were mapped from the SuCas12a2 binary crystal structure (PDB 8D49) [9]. Amino acid conservation at key functional residues was scored as identical, conservative (preserves the chemical property), or non-conservative (major chemical change) relative to SuCas12a2.

## Supplementary Tables

**Table S1.**
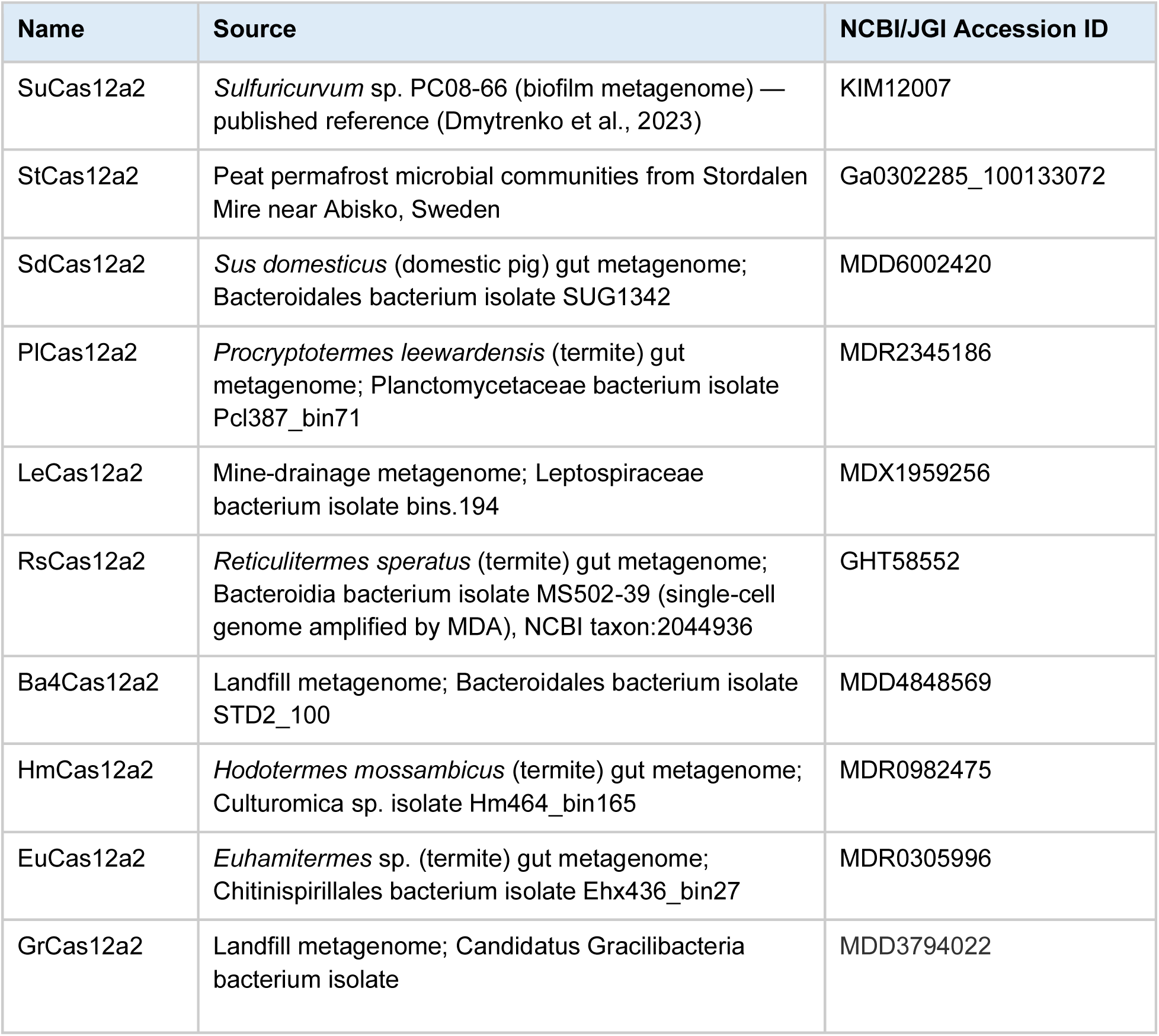
Cas12a2 ortholog panel.

---

**Supplementary Figure 1.**
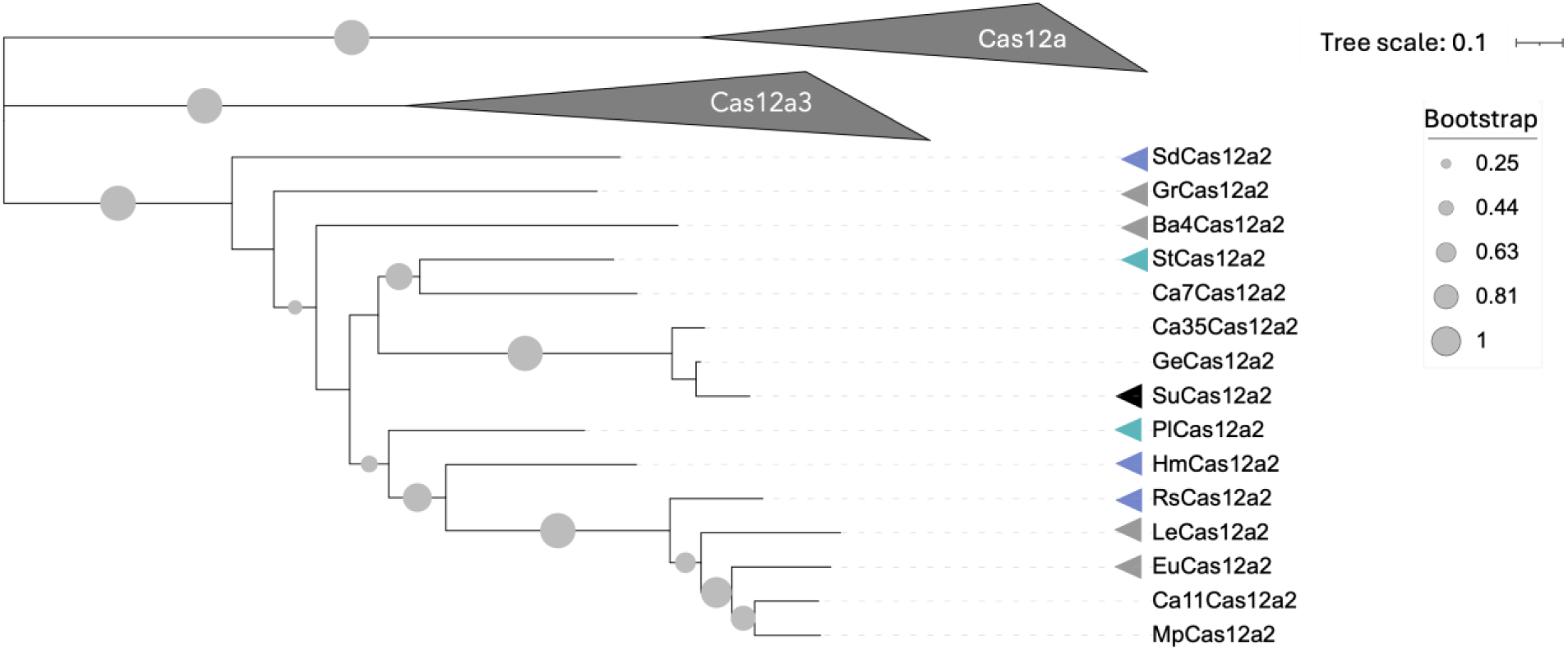
Maximum likelihood phylogenetic tree of 24 Cas12 nuclease, with outgroup Cas12a and Cas12a3 clades collapsed. Coloring corresponds to subsequent activity designations based on functional characterization. The tree was calculated using MEGA12 [16] from an amino acid alignment (MUSCLE [18]) that was trimmed using ClipKIT [17]. The tree was visualized using iTOL [19]. Bootstrap support for the major nodes are indicated. The scale bar indicates evolutionary distance based on substitution rate.

**Supplementary Figure 2.**
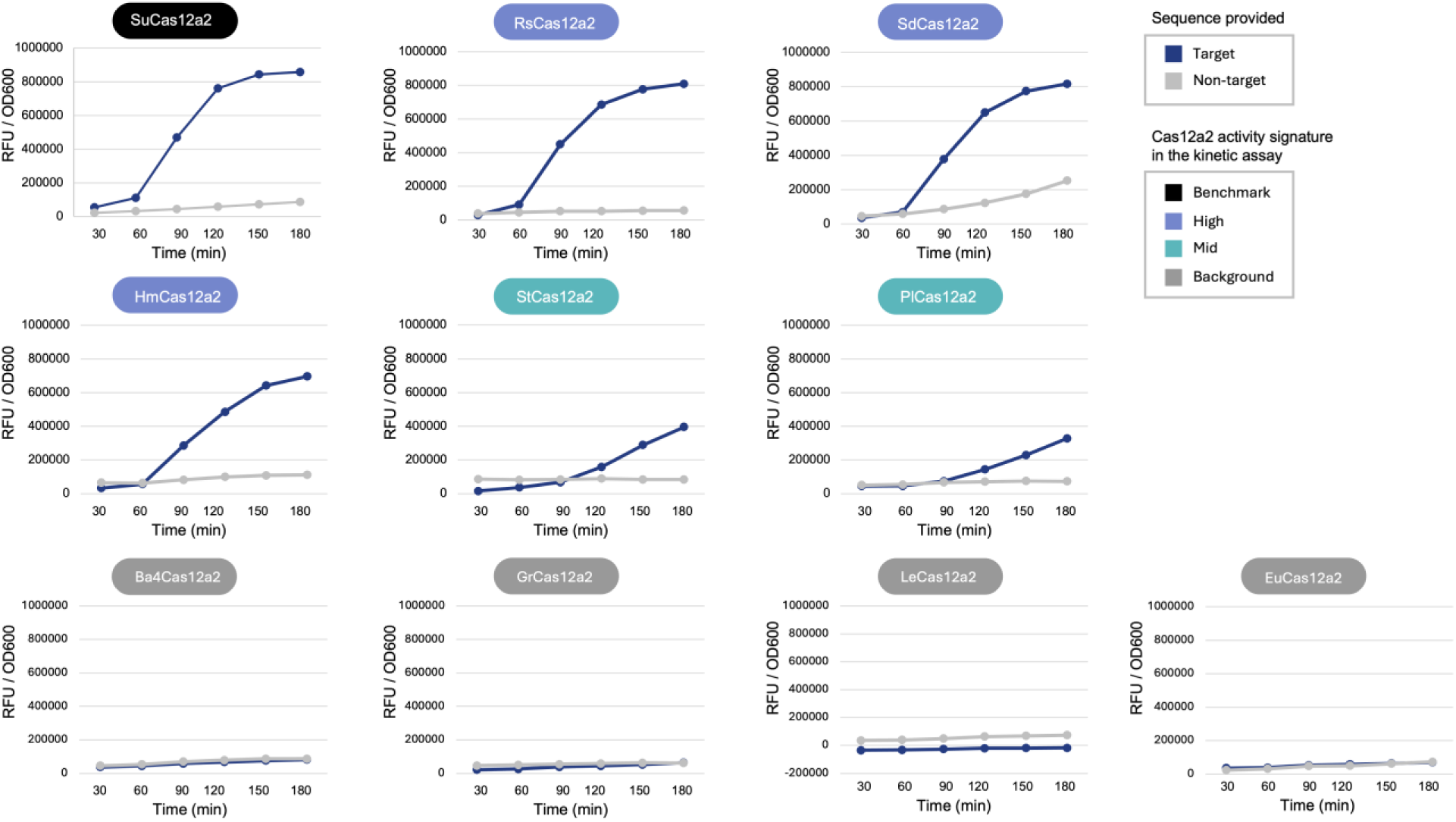
Cas12a2 orthologs have different signatures of activity in the DNA damage kinetic assay. Representative kinetic traces of *recA* driven GFP signal (RFU/OD600) over time for Cas12a2 orthologs against target (blue) and non-target (grey) plasmids.

**Supplementary Figure 3.**
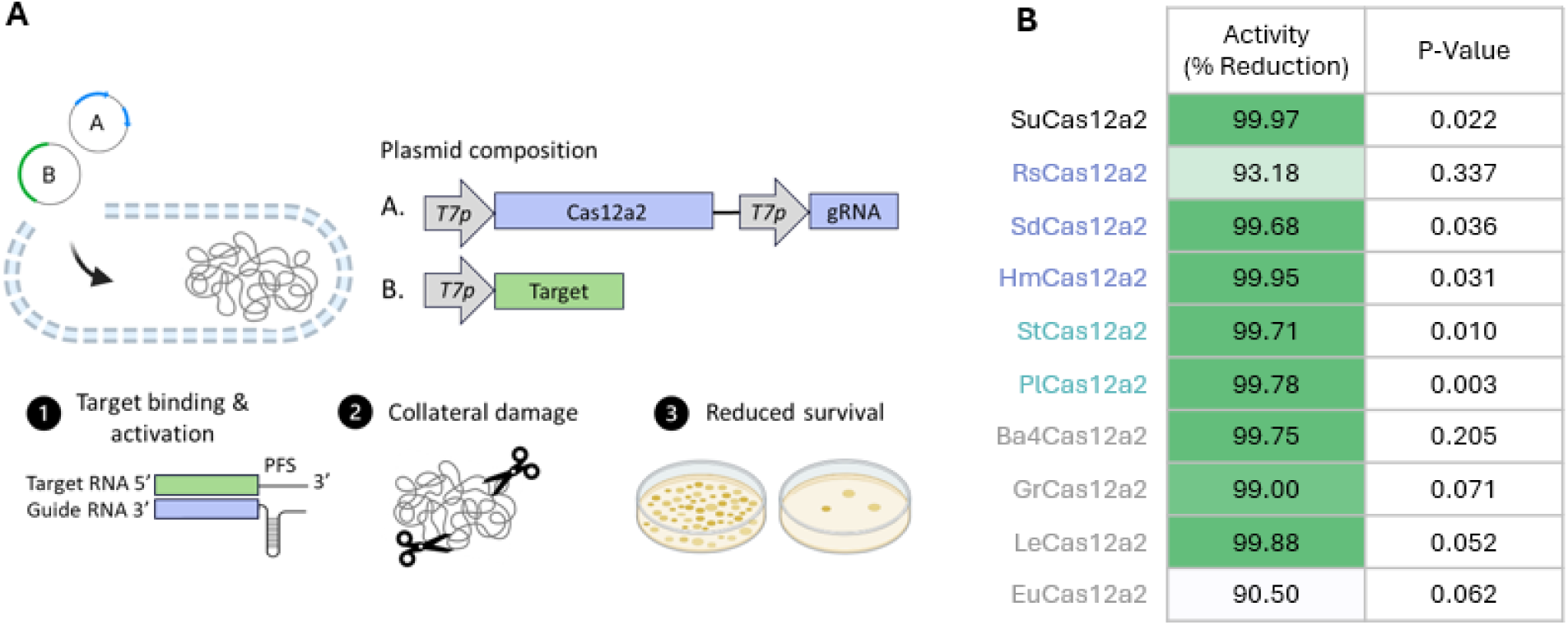
(A) Schematic of the plate-based bacterial toxicity assay. Strains were transformed with two plasmids carrying inducible Cas12a2 and gRNA (plasmid A) and inducible target sequence (plasmid B). Binding of target RNA activates Cas12a2 collateral activity resulting in reduced colony survival. (B) Preliminary screening results for 9 new nucleases classified as functional based on % reduction of surviving colonies in the toxicity assay.

**Supplementary Figure 4.**
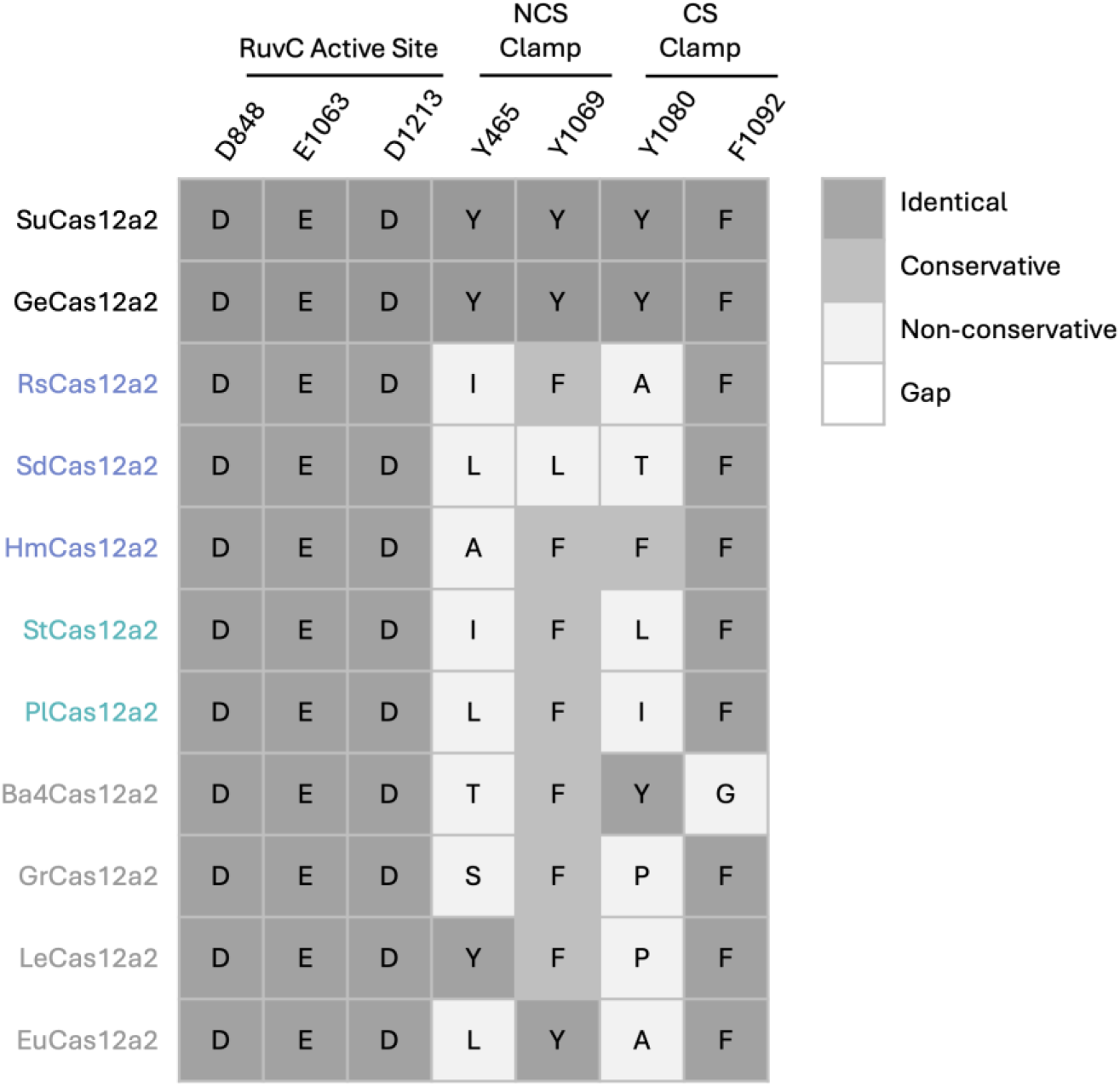
Newly characterized Cas12a2 nucleases have variation at key functional residues. Key residue mapping of the two previously characterized Cas12a2 nucleases (SuCas12a2 [8,9] and GeCas12a2 [12]) and newly characterized Cas12a2 nucleases color-coded based on activity designation in the DNA damage kinetic assay. Amino acid conservation at each site was scored as identical, conservative (preserves the chemical property), or non-conservative (major chemical change) relative to SuCas12as2.

## Supplementary Sequences

Supplementary sequences for Cas12a outgroups (4 sequences), Cas12a2 orthologs (15 sequences), and Cas12a3 outgroups (5 sequences) are provided below in FASTA format.

### Cas12a outgroups

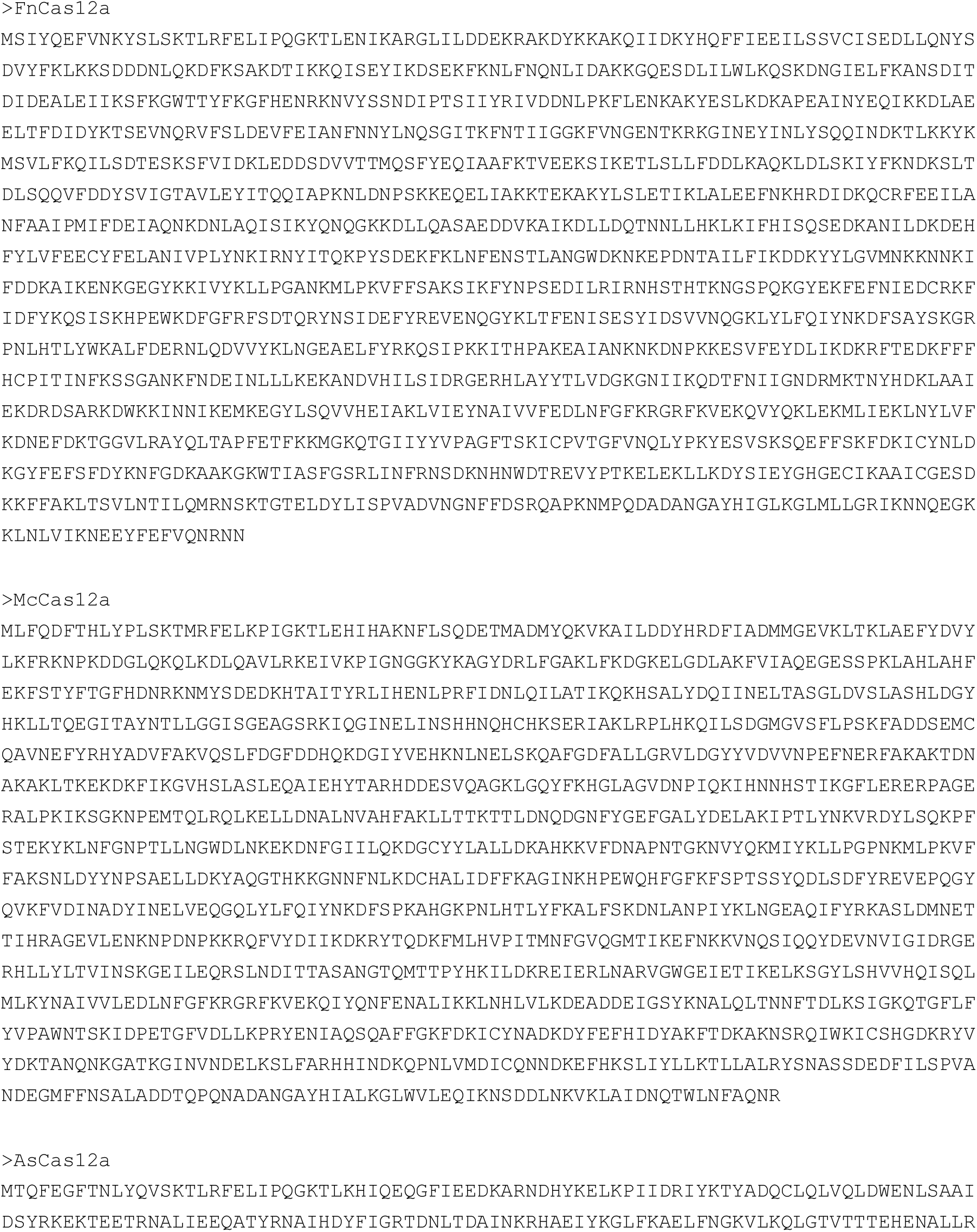

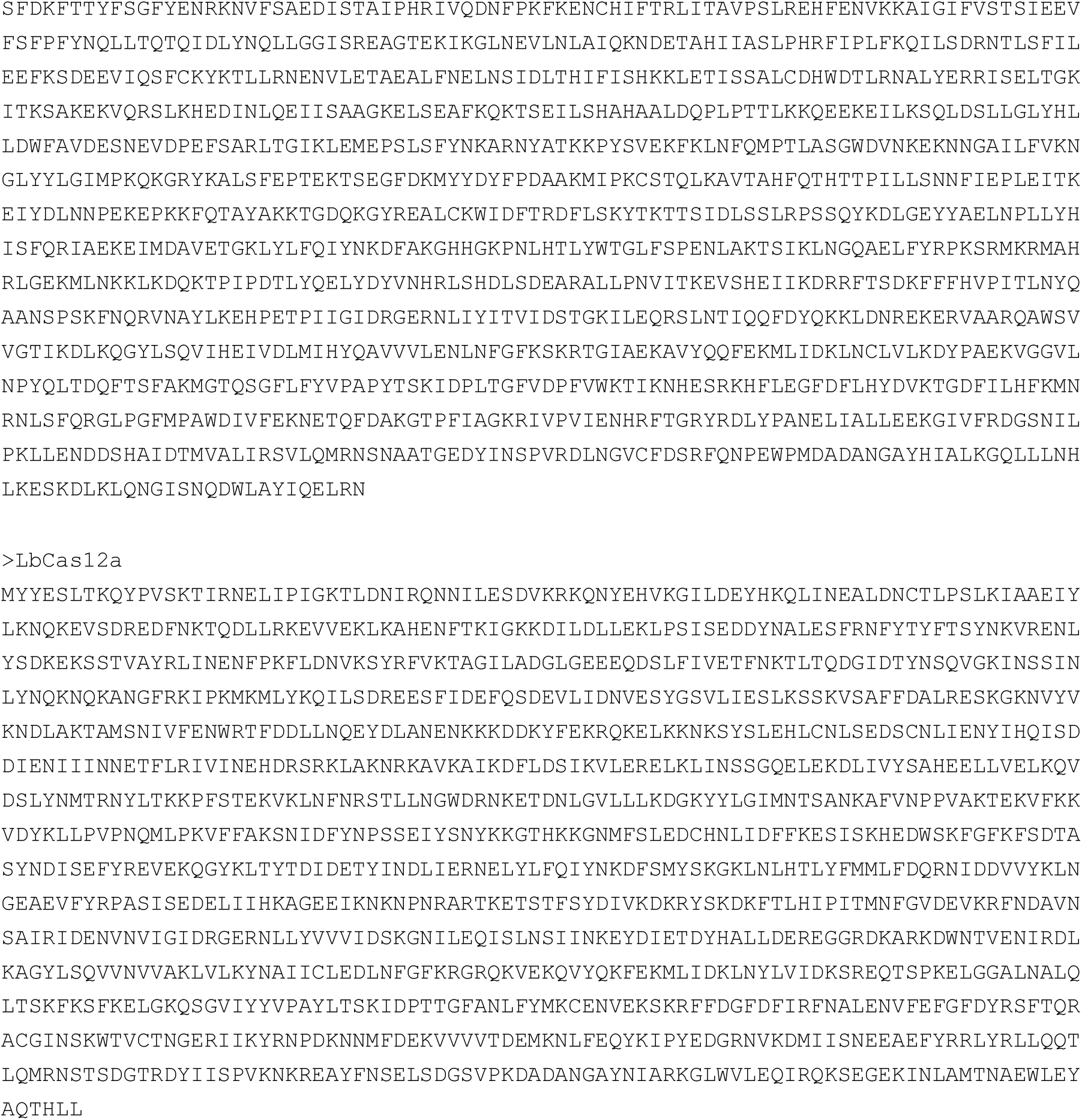

### Cas12a2 orthologs

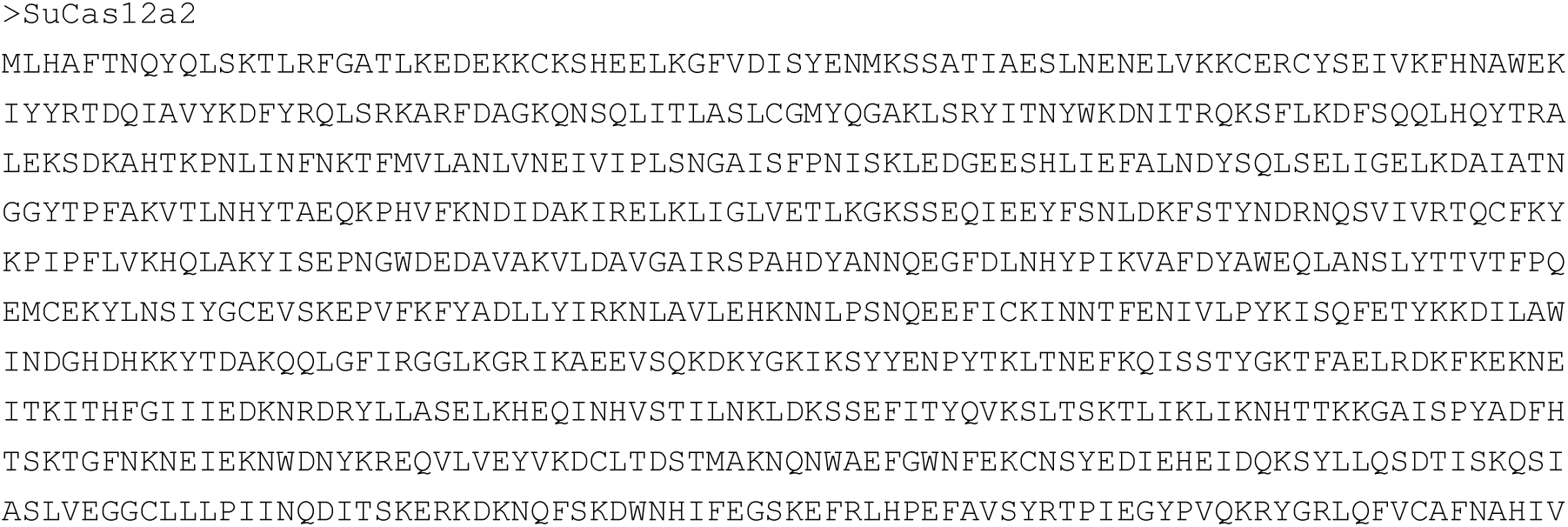

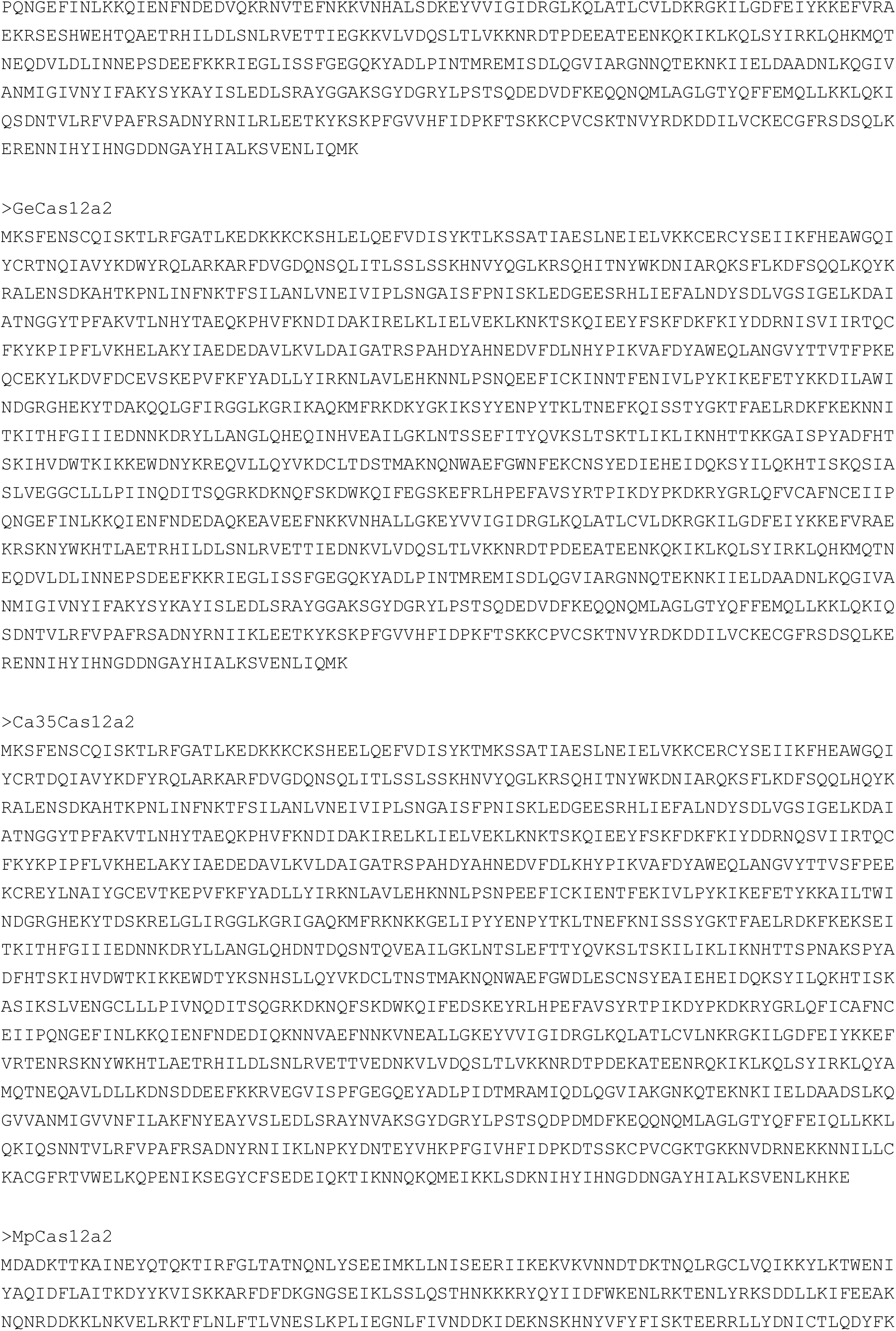

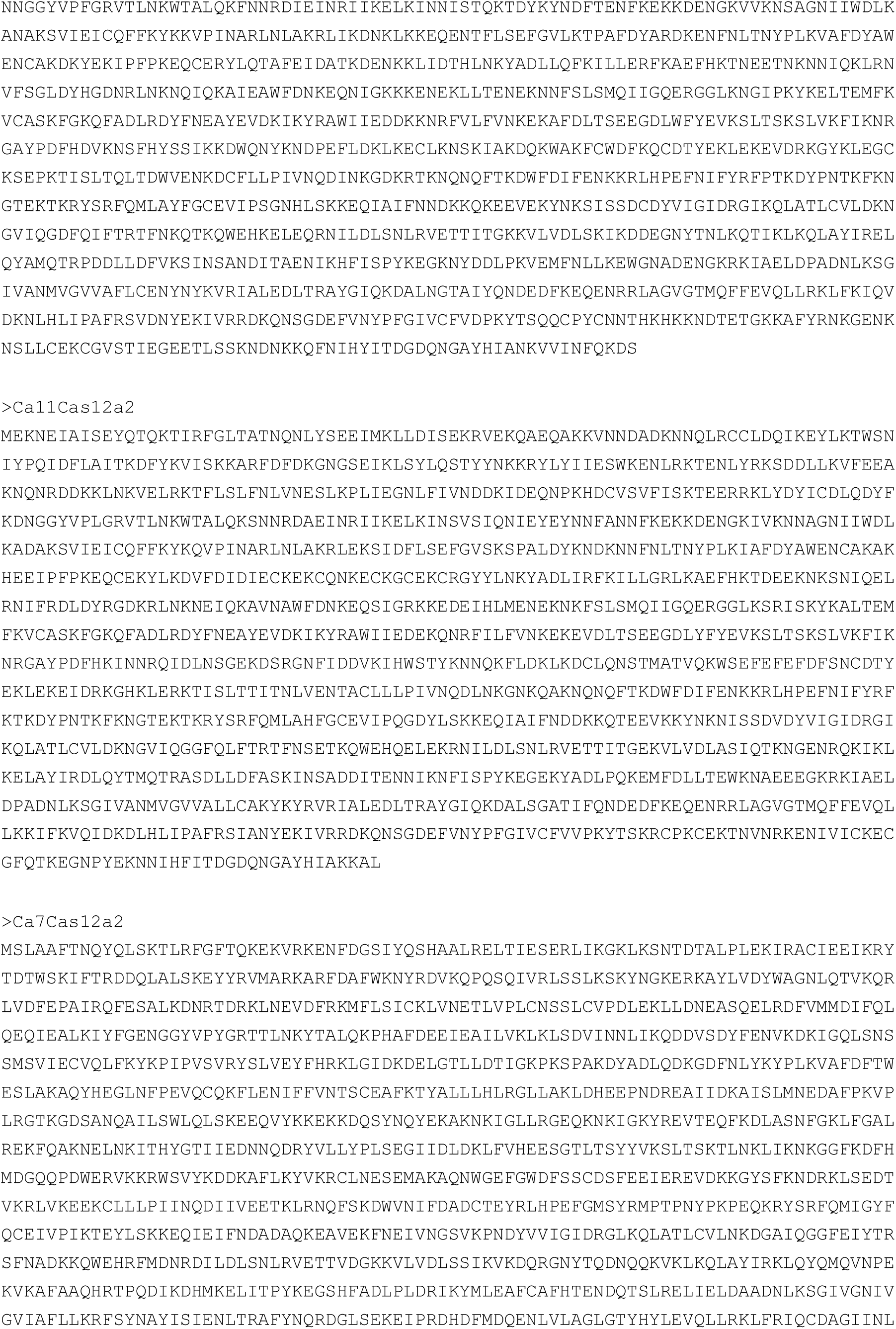

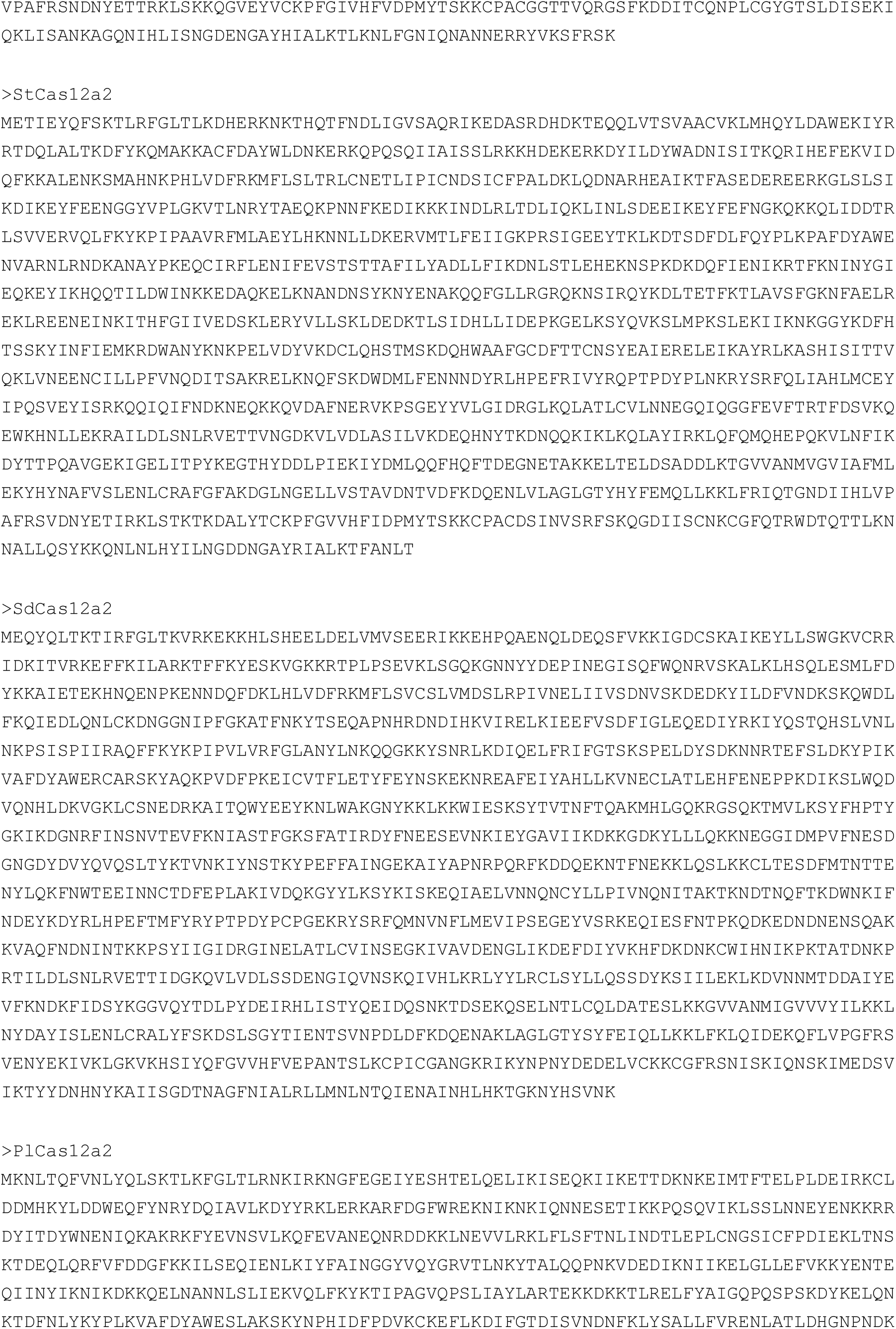

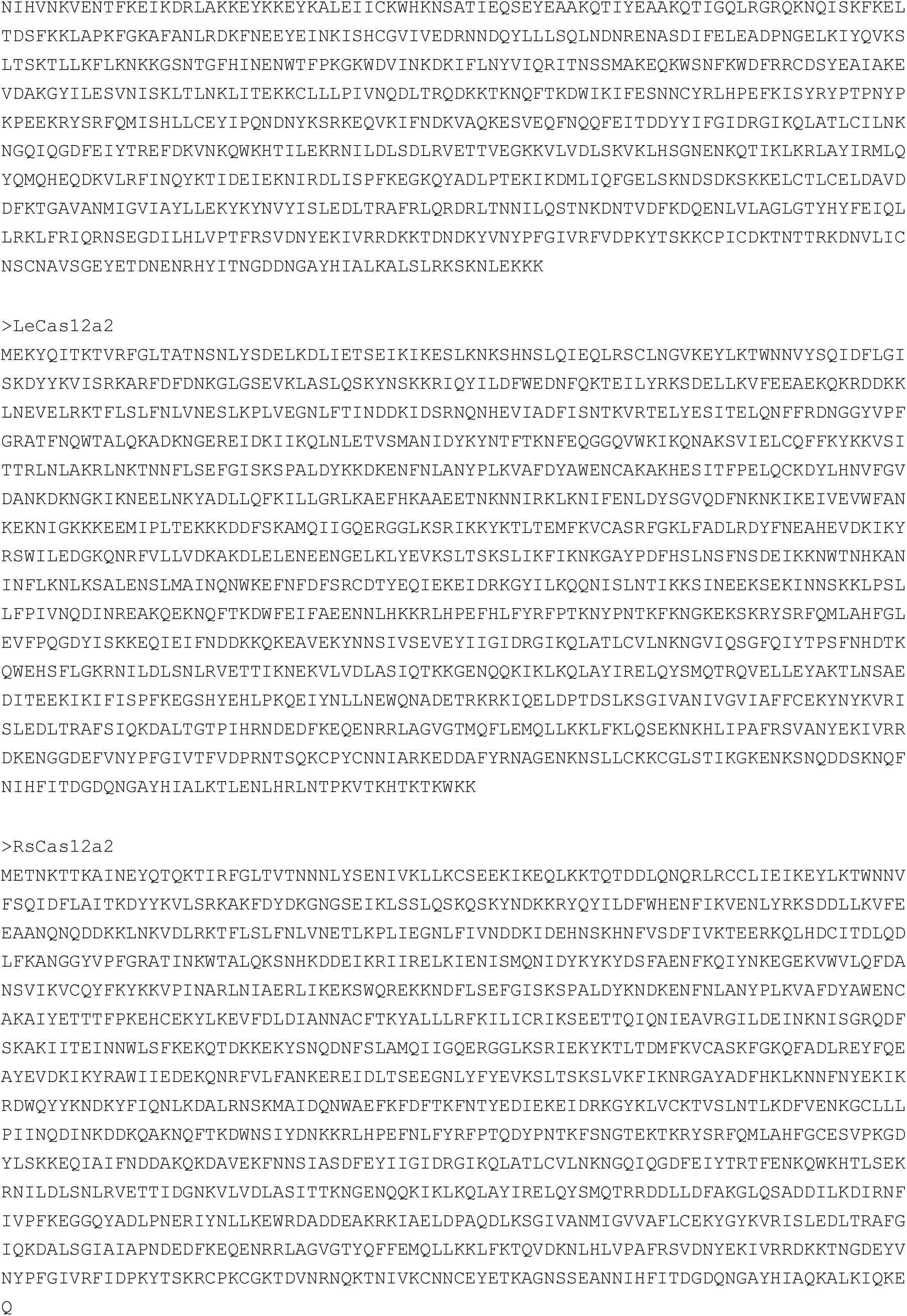

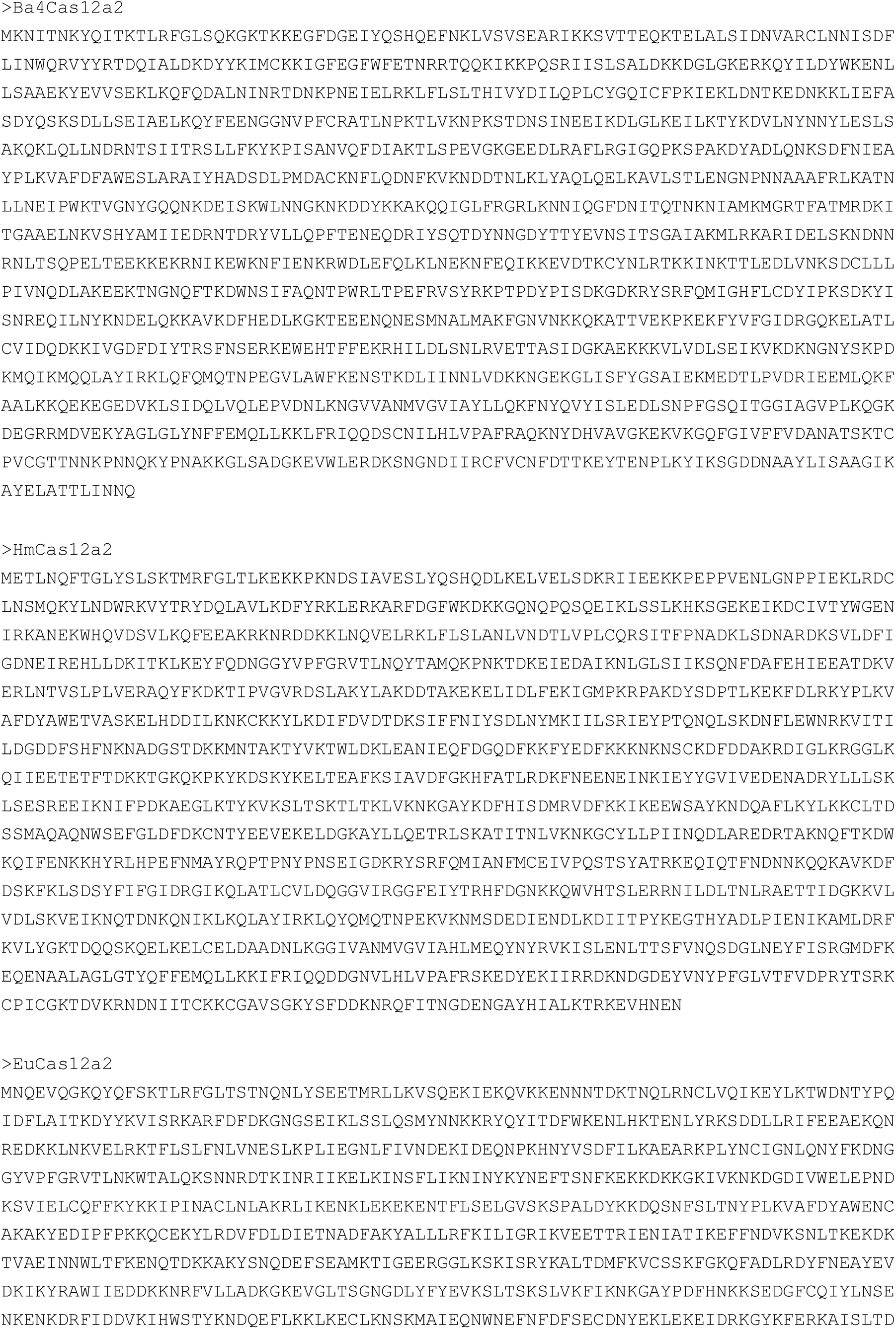

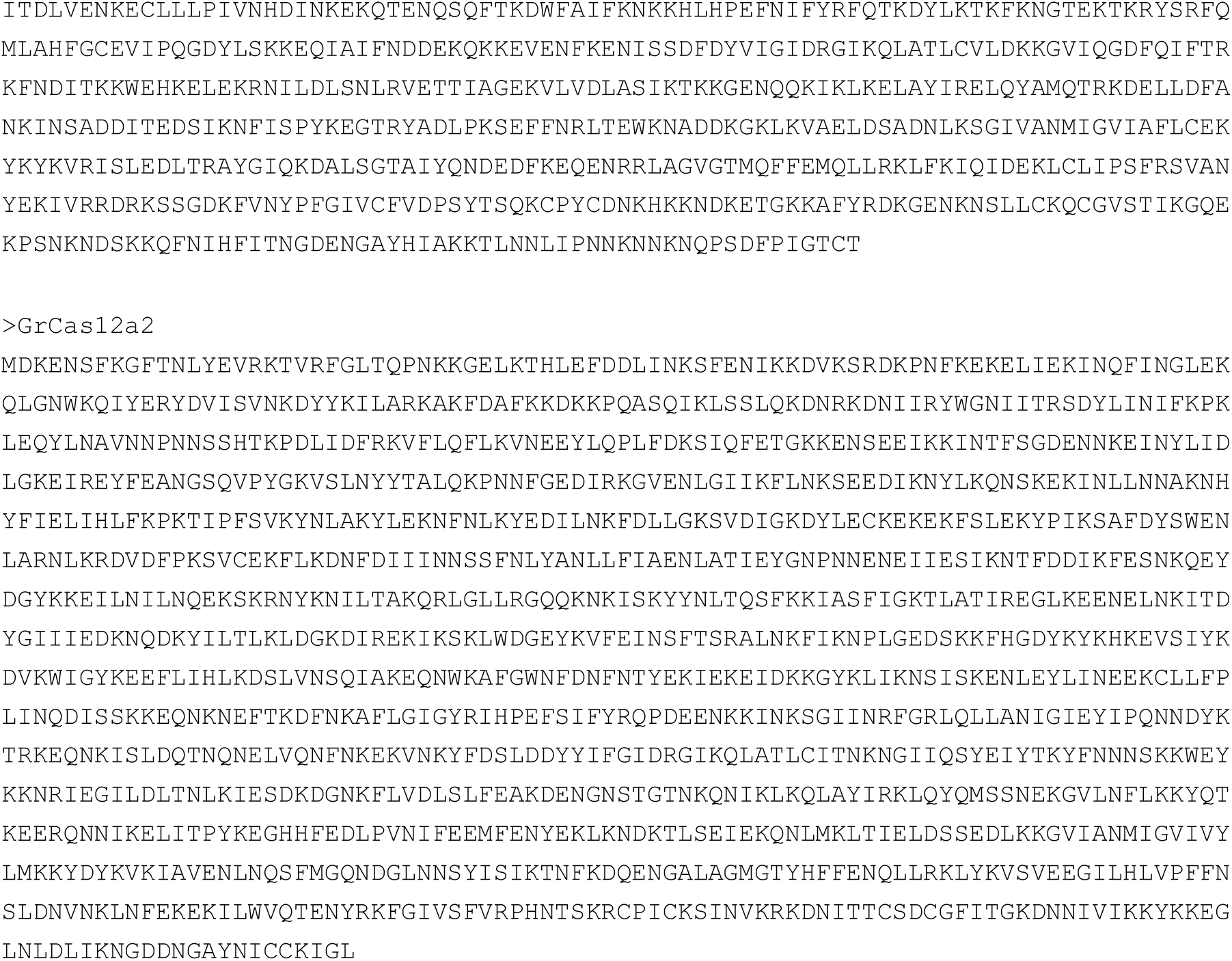

### Cas12a3 outgroups

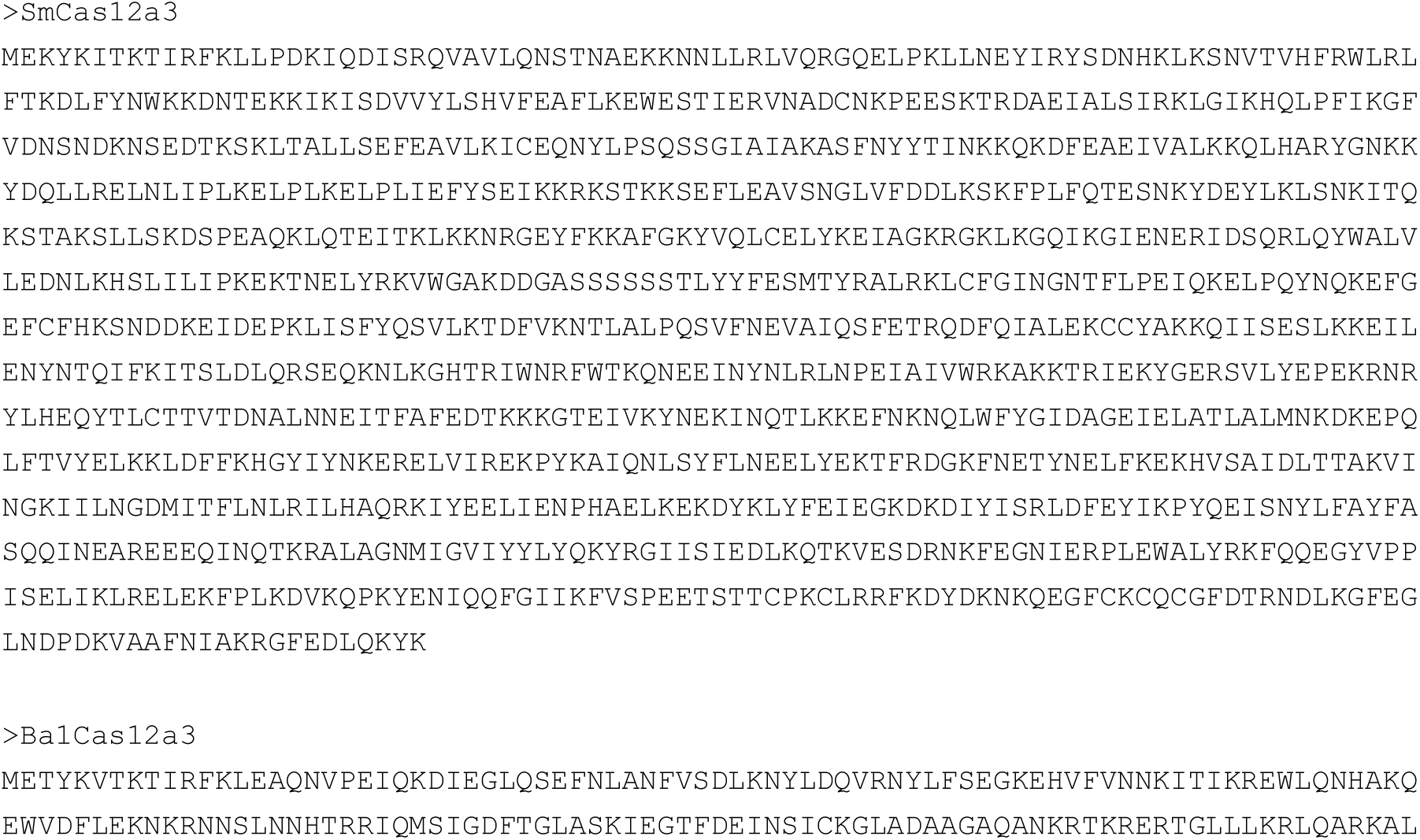

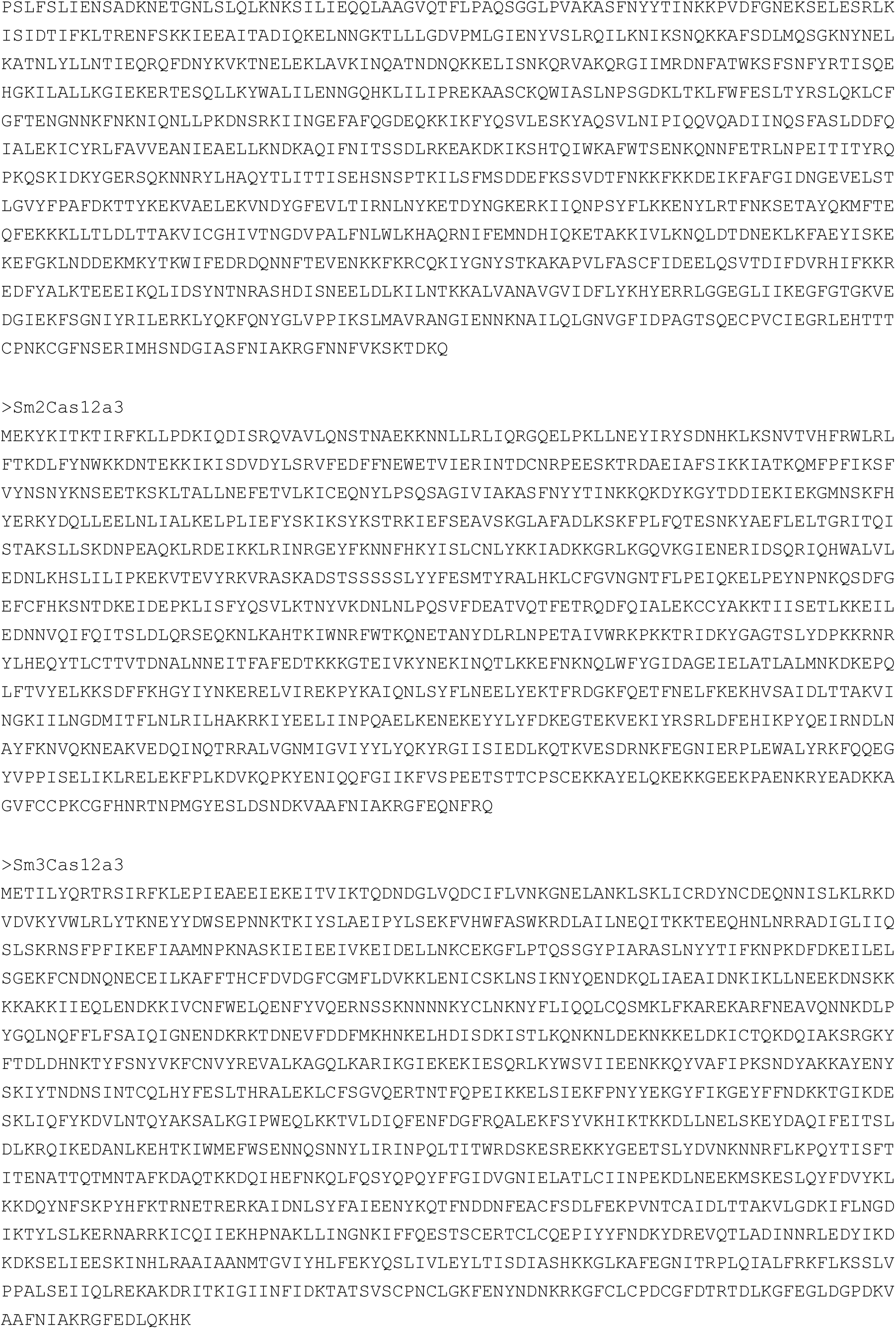

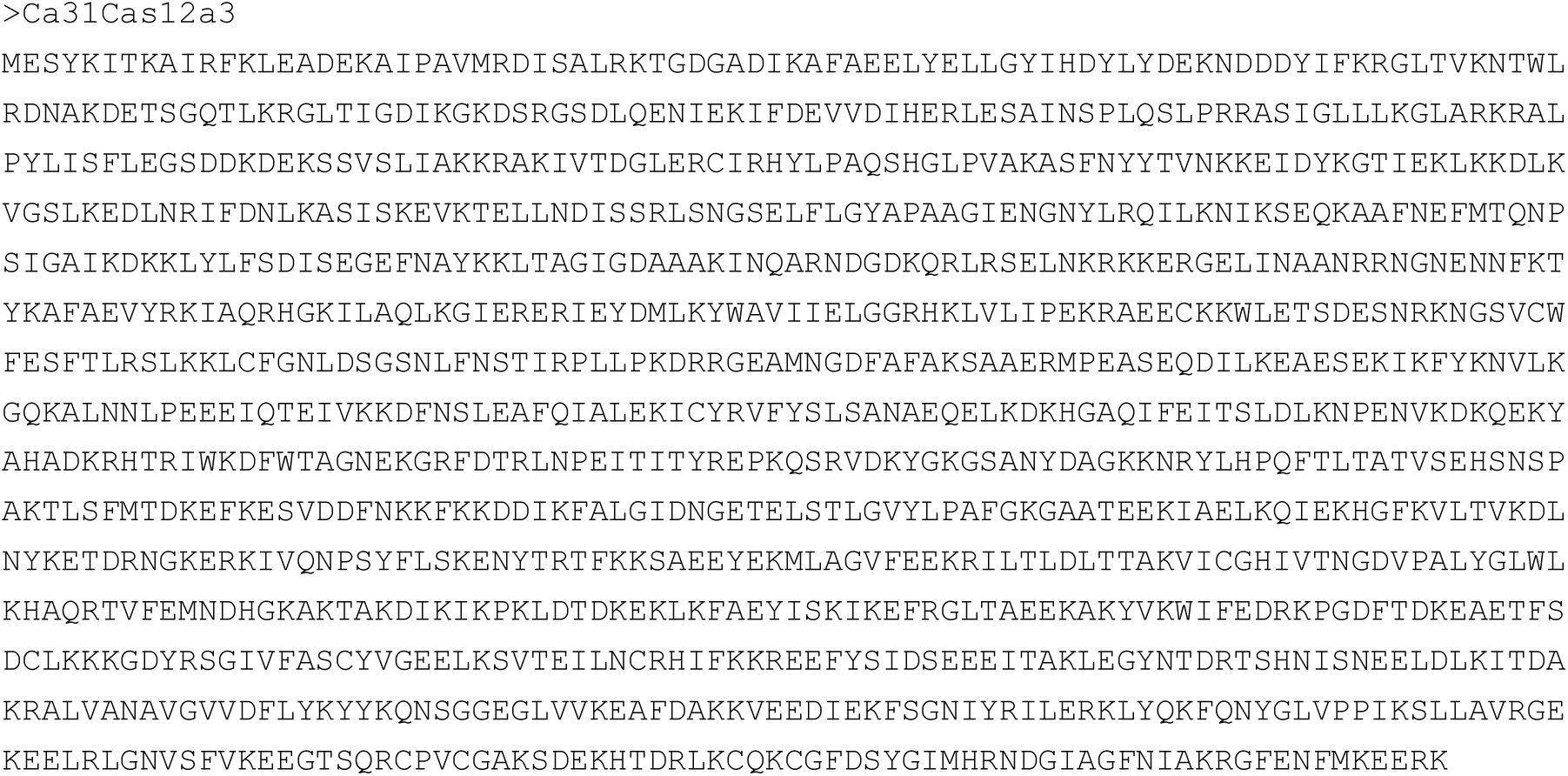

